# Functional genomics identifies AMPD2 as a new prognostic marker for undifferentiated pleomorphic sarcoma

**DOI:** 10.1101/292805

**Authors:** Martin F. Orth, Julia S. Gerke, Thomas Knösel, Annelore Altendorf-Hofmann, Julian Musa, Rebeca Alba Rubio, Stefanie Stein, Marlene Dallmayer, Michaela C. Baldauf, Aruna Marchetto, Giuseppina Sannino, Shunya Ohmura, Jing Li, Michiyuki Hakozaki, Thomas Kirchner, Thomas Dandekar, Elke Butt, Thomas G. P. Grünewald

**Author notes:** **correspondence** Thomas G. P. Grünewald, M.D., Ph.D. Max-Eder Research Group for Pediatric Sarcoma Biology Institute of Pathology, Faculty of Medicine, LMU Munich Thalkirchner Str. 36, 80337 Munich, Germany Web www.lmu.de/sarkombiologie.

## Abstract

Soft-tissue sarcomas are rare, heterogeneous and often aggressive mesenchymal cancers. Many of them are associated with poor outcome, in part because biomarkers that can reliably identify high-risk patients are lacking. Studies on sarcomas often are limited by small sample sizes rendering the identification of novel biomarkers difficult when focusing only on individual cohorts. However, the increasing number of publicly available ‘omics’ data opens inroads to overcome this obstacle.

Here, we combine high-throughput transcriptome analyses, immunohistochemistry, and functional assays to show that high adenosine monophosphate deaminase 2 (*AMPD2*) is a robust prognostic biomarker for worse patient outcome in undifferentiated pleomorphic sarcoma (UPS). Publicly available gene expression and survival data for UPS from two independent studies, The Cancer Genome Atlas (TCGA) and the CINSARC reference dataset, were subjected to survival association testing. Genes, whose high expression was significantly correlated with worse outcome in both cohorts (overall and metastasis-free survival), were considered as prognostic marker candidates. The best candidate, *AMPD2*, was validated on protein level in an independent tissue microarray. Analysis of DNA copy-number and matched gene expression data indicated that high *AMPD2* expression is significantly correlated with copy-number gains at the *AMPD2* locus. Gene-set enrichment analyses of *AMPD2* co-expressed genes in both UPS gene expression datasets suggested that highly *AMPD2* expressing tumors are enriched in gene signatures involved in tumorigenesis. Consistent with this prediction in primary tumors, knockdown of *AMPD2* by RNA interference with pooled siRNAs or a doxycycline-inducible shRNA construct in the UPS cell line FPS-1 markedly inhibited proliferation *in vitro* and tumorigenicity *in vivo*.

Collectively, these results provide evidence that *AMPD2* may serve as a novel biomarker for outcome prediction in UPS. Our study exemplifies how the integration of available ‘omics’ data, immunohistochemical analyses, and functional experiments can identify novel biomarkers even in a rare sarcoma, which may serve as a blueprint for biomarker identification for other rare cancers.

## INTRODUCTION

Soft-tissue sarcomas comprise a broad range of mesenchymal tumors^1^, which are often characterized by high relapse and metastasis rates^2,3^. For many sarcoma entities there are presently no prognostic biomarkers available that could help in tailoring individualized therapy^4^. Therefore, most patients are treated with radical and sometimes mutilating surgical resections, combined with various chemotherapeutic and/or irradiation protocols^2,3^.

Despite the general rarity of sarcomas, The Cancer Genome Atlas (TCGA) consortium provides a multidimensional genetic and clinically annotated dataset for various rare tumor entities including undifferentiated pleomorphic sarcoma (UPS)^5^, which constitutes a major clinical challenge^6^.

The diagnosis of UPS is established for high-grade malignant neoplasms characterized by tumor cells with diffuse pleomorphism and the absence of a specific line of differentiation^1^. Most UPS occur in the extremities at an age of over 60 years^7^. Although these sarcomas are heterogeneous, they are commonly aggressive. While the 5-year overall survival is around 60%, metastasis-free survival in the same time interval is only around 30%^8^. In fact, UPS tend to frequently metastasize, especially to the lungs (40-50%)^1^. The standard treatment for UPS comprises limb sparing resection combined with either neoadjuvant and/or adjuvant radiotherapy^9^. So far, no clear benefit of other and more aggressive therapy regimens has been proven^7,9^, possibly due to the lack of prognostic markers that can discriminate high-risk from low-risk patients.

In this study, we combined bioinformatic analyses of the TCGA and an additional gene expression dataset for UPS, immunohistochemical analysis of a tissue microarray (TMA), and *in vitro* and *in vivo* experiments to probe for robust prognostic markers in UPS. We found that high expression of the adenosine monophosphate deaminase 2 (*AMPD2*), which is involved in purine metabolism^10^, is associated in all independent cohorts with worse patient outcome, possibly by promoting proliferation of UPS cells.

## MATERIALS AND METHODS

### Retrieval and processing of gene expression and clinical data

Normalized mRNA expression levels from RNA sequencing (RNA-seq) data of UPS were retrieved together with corresponding clinical annotations and further normalized genetic readouts (copy number variation) from the TCGA data portal. RNA expression data from the CINSARC (complexity index in sarcomas) microarray analysis of UPS (GSE21050, Affymetrix HG-U133-Plus2.0 microarray)^11^ were downloaded from the Gene Expression Omnibus (GEO). The corresponding clinical data for GSE21050 were extracted from the attached series matrix file. All CEL files were manually inspected to be generated on the same microarray and simultaneously normalized with Robust Multi-array Average (RMA)^12^ including background correction, between-array normalization, and signal summarization using a custom brainarray chip description file (CDF; v19 ENTREZG), yielding one optimized probe set per gene^13^. Patient characteristics for both cohorts are given in Table 1.

**Table 1.** Characteristics of the UPS cohorts

| **Feature** | **TCGA** | **GSE21050** | **TMA** |
| --- | --- | --- | --- |
| *n* of samples | 50 | 129 | 83 |
| Expression level assessed | RNA | RNA | Protein |
| Mode of assessment | RNASeqV2 | Microarray (Affymetrix) | Immunoreactivity scoring |
| Age range (median) [years] | 42-90 (68) | NA | 19-79 (56) |
| Event type | Overall survival | Metastasis-free survival | Overall survival |
| *n* event (%) | 12 (24) | 39 (30) | 36 (43) |
| Follow-up time range (median) [month] | 1.1-124.4 (20.61) | 0.36-211.8 (32.8) | 2-288 (150) |

### Analysis of association with patient outcome in gene expression datasets

For both RNA expression datasets patients were stratified in two subgroups defined by either high or low expression of a given gene (cut-off: median expression level of the given gene). Differential event-free survival in both subgroups was assessed in each cohort independently with the Mantel-Haenszel test, a time-stratified modification of the log-rank test. Events were defined in the TCGA cohort as ‘death’, and in the GSE21050 cohort as ‘occurrence of metastasis’. This analysis was performed automatically for all 17,536 genes represented in both expression datasets with an in-house software. Those genes whose high expression showed a concordant and significant (*P* < 0.01; without correction for multiple testing) association with worse event-free survival in both cohorts, were manually inspected and displayed using the Kaplan-Meier method in the GraphPad Prism 5 software.

### Linear regression analysis for copy-number variations (CNVs) and gene expression

To evaluate if *AMPD2* expression is regulated by CNVs in UPS, the log2 transformed segment mean values for the copy-number at the *AMPD2* locus and the *AMPD2* expression of the TCGA UPS cohort were displayed and subjected to linear regression analysis in GraphPad Prism 5 software.

### Gene Set Enrichment Analysis (GSEA)

To identify gene sets that are enriched among *AMPD2*-co-regulated genes, all genes in both gene expression datasets were ranked by their Pearson correlation coefficient with *AMPD2* expression and a pre-ranked GSEA with 1,000 permutations was performed^14^.

### Immunohistochemistry and immunoreactivity scoring

A tissue microarray (TMA) provided by T. Knösel comprising 83 UPS, each represented by two cores (1 mm in diameter), was used to analyze the correlation of AMPD2 expression levels with overall survival. Antigen retrieval was carried out by heat treatment with ProTaqs IV Antigen Enhancer (401602392, Quartett, Berlin, Germany). Next, the TMA slides were stained with a polyclonal anti-AMPD2 antibody raised in rabbit (1:1,300; HPA050590, Atlas Antibodies, Bromma, Sweden) for 60 min at RT, followed by a monoclonal secondary horseradish peroxidase (HRP)-coupled horse-anti-rabbit antibody (ImmPRESS Reagent Kit, MP-7401, Vector Laboratories, Burlingame, USA). AEC-Plus (K3469, Agilent, Santa Clara, USA) was used as chromogen. Samples were counterstained with hematoxylin (H-3401, Vector Laboratories). The specificity of the used anti-AMPD2 antibody was validated by staining of xenografts from an UPS cell line (FPS-1) with an experimentally induced knockdown of AMPD2 (Fig. 4). In the TMA, the AMPD2 immunoreactivity was scored by a consultant pathologist (T. Knösel), who was blinded to the clinical data. Cytosolic AMPD2 immunoreactivity was classified into ‘no’, ‘low’, ‘intermediate’, and ‘strong’. Assignment of the scoring results to the clinical data was performed by an independent statistician (H. Altendorf-Hofmann). Association with overall survival was calculated with the Kaplan-Meier method and Mantel-Haenszel test in SPSS.

Formalin-fixed and paraffin-embedded (FFPE) xenografts of the FPS-1 UPS cell line were stained with hematoxylin and eosin (H&E). AMPD2 staining was carried out as described above. For cleaved caspase-3 staining, antigen retrieval was carried out by heat treatment with Target Retrieval Solution Citrate pH6 (S2369, Agilent). Then, slides were incubated with the polyclonal cleaved caspase-3 primary antibody raised in rabbit (1:100; 9661, Cell Signaling, Frankfurt am Main, Germany) for 60 min at RT followed by ImmPRESS Reagent Kit. DAB+ (K3468, Agilent) was used as chromogen, hematoxylin for counterstaining. Ki-67staining was performed with a VENTANA BenchMark system (Roche, Basel, Switzerland) with an ultraView detection kit (Roche) and a monoclonal mouse anti-Ki-67 antibody (M7240, Agilent).

### Cell lines and cell culture conditions

The FPS-1 UPS cell line was established and described previously^15^. HEK293T cells were obtained from ATCC (Manassas, USA). Both FPS-1 and HEK293T cells were grown in RPMI 1640 medium (Merck, Darmstadt, Germany), supplemented with 15% or 10% tetracycline-free fetal bovine serum (Biochrom, Berlin, Germany), respectively, and 100 U/ml penicillin and 100 µg/ml streptomycin (Merck) in a humidified 5% CO_2_ atmosphere at 37°C. Cells were subcultured every two to seven days, detaching the cells with trypsin/EDTA (Merck). Cells were only used up to 15 passages, checked routinely for the absence of mycoplasma by nested PCR, and maintenance of cell line identity was verified by STR-profiling.

### RNA interference (RNAi)

Transient suppression of *AMPD2* expression was achieved by transfection of FPS-1 cells with 10 nM of an siPOOL (siTOOLs, Planegg, Germany) consisting of 30 different short interfering RNAs (siRNAs) targeting *AMPD2* (sipAMPD2), which virtually eliminates off-target effects^16^. A commercial non-targeting siPOOL (siTOOLs) was used as a control (sipControl). The siPOOLs were complexed with the transfection reagent HiPerfect (Qiagen, Venlo, The Netherlands) and target cells were reversely transfected. After 24 h the transfection reagent was diluted by doubling the amount of cell culture medium. Knockdown efficacy was validated by quantitative real-time polymerase chain reaction (qRT-PCR).

### Generation of FPS-1 cells with an inducible *AMPD2* targeting shRNA

To study the effects of *AMPD2* knockdown for a longer period of time, we employed the lentiviral pLKO-TET-ON all-in-one vector system containing a puromycin selection cassette^17^. Herein, we cloned either a non-targeting short hairpin RNA (shControl; 5’-CAACAAGATGAAGAGCACCAA-3’) or an shRNA against *AMPD2* (shAMPD2; target sequence 5’-GGGTATCTGGGAAGTACTTTG-3’). Vectors were expanded in Stellar Competent Cells (Clontech, Kyoto, Japan) and clones with integrated shRNA were sequenced by Sanger sequencing to validate the correct shRNA sequence. Lentivirus was produced in HEK293T cells, which were transfected using the Lipofectamine LTX Plus Reagent system (Life Technologies, Darmstadt, Germany). Target cells were infected with filtered supernatant of the HEK293T cells without Polybrene. Before the cells reached confluence, successfully transduced cells were selected with 1 µg/ml puromycin (InvivoGen, San Diego, USA). For induction of shRNA expression, 0.5 µg/ml doxycycline (dox) (Merck) was added to the culture medium.

### Analysis of cell proliferation

For assessment of proliferation, 4 × 10^5^ FPS-1 cells were seeded in wells of 6-well plates and transfected with either sipAMPD2 or sipControl. Medium was doubled after 24 h. 48 h after transfection cells were re-transfected. Cells including their supernatant were counted 120 h after initial transfection using the Trypan-Blue (Sigma-Aldrich, Taufkirchen, Germany) exclusion method and standardized hemocytometers (C-Chip, Biochrom). The same assays were carried out with dox-inducible shRNA FPS-1 infectants, in which the *AMPD2* knockdown was induced by addition of 0.5 µg/ml dox after 24 h. Doxycycline was refreshed after 72 h.

### Analysis of tumor growth *in vivo*

2 × 10^6^ FPS-1 cells transduced with inducible shRNA against *AMPD2* were injected in the right flank of NOD/Scid/gamma (NSG) mice. Tumor growth was measured every second day with a caliper. When most tumors reached an average diameter of 5 mm, mice were randomized into two groups of which one was treated henceforth with 2 mg/ml dox (bela-pharm, Vechta, Germany) dissolved in drinking water containing 5% sucrose (Sigma-Aldrich), and the other one with 5% sucrose only. Mice were sacrificed when the average tumor diameter exceeded 15 mm (stop criterion). Tumors were quickly extracted, small samples were snap frozen in liquid nitrogen prior to RNA isolation, and the remaining tumor tissue was formalin-fixed (4%) and paraffin-embedded for (immuno)histology. Animal experiments were carried out in accordance with recommendations of the European Community (86/609/EEC), local authorities, and the UKCCCR guidelines (guidelines for the welfare and use of animals in cancer research).

### RNA extraction, reverse transcription, and qRT-PCR

Total RNA was extracted using the NucleoSpin RNA kit (Macherey-Nagel, Düren, Germany) according to the manufacturer’s protocol. RNA (1 µg) was reversely transcribed with the High Capacity cDNA Reverse Transcription Kit (Thermo Fisher Scientific, Rockford, USA), according to the manufacturer’s protocol. For qRT-PCR 1:10 diluted cDNA and 0.5 µM forward and reverse primer for *AMPD2* and the housekeeping gene *RPLP0* were used in SYBR Select Master Mix (Thermo Fisher Scientific) (total reaction volume: 15 µl) in a Bio-Rad CFX Connect Real-Time PCR Detection System (Bio-Rad, Munich, Germany). All primers were purchased from Eurofins Genomics (Ebersberg, Germany). Primer sequences were as follows:

*AMPD2* forward 5’-GGTCTCTGCATGTCTCCATTC-3’;

*AMPD2* reverse 5’-CTCAATACCTGGGCCATCAG-3’;

*RPLP0* forward 5’-GAAACTCTGCATTCTCGCTTC-3’;

*RPLP0* reverse 5’-GGTGTAATCCGTCTCCACAG-3’.

The qRT-PCR was carried out with the following thermal conditions: heat activation at 95°C for 2 min, DNA denaturation at 95°C for 10 sec, and annealing and elongation at 60°C for 20 sec (50 cycles), final denaturation at 95°C for 30 sec. *AMPD2* expression before and after knockdown was calculated with the Delta-Delta-Cq method.

## RESULTS

### High *AMPD2* mRNA levels correlate with worse outcome in two independent UPS cohorts

To identify prognostic marker candidates for UPS, we stepwise crossed two independent datasets for which matched clinical data and transcriptome-wide expression profiles were available (Fig. 1a). The first was derived from The Cancer Genome Atlas (TCGA) sarcoma panel (*n* = 50)^5^, the second from the CINSARC reference dataset (*n* = 129; GSE21050)^11^. We calculated independently for both datasets for each gene (17,536 genes represented in both cohorts) the association with event-free survival (TCGA: event = death; GSE21050: event = occurrence of metastasis) when stratifying each dataset by the median expression of the given gene (see methods section). This yielded 379 genes in the TCGA and 682 genes in the GSE21050 cohort with a *P* value of < 0.01 (**Supplementary Tables 1 and 2**). Notably, only five genes were significantly (*P* < 0.01) and concordantly positively associated with worse event-free survival in both cohorts, namely *AMPD2*, *CA2, NAV2*, *RGS3*, *SLC35D1* (Fig. 1b). Interestingly, we noted that for *AMPD2* the survival curves in the Kaplan-Meier analyses separated very early, which could indicate a potential biological relevance (Fig. 1c).

**Figure 1.**
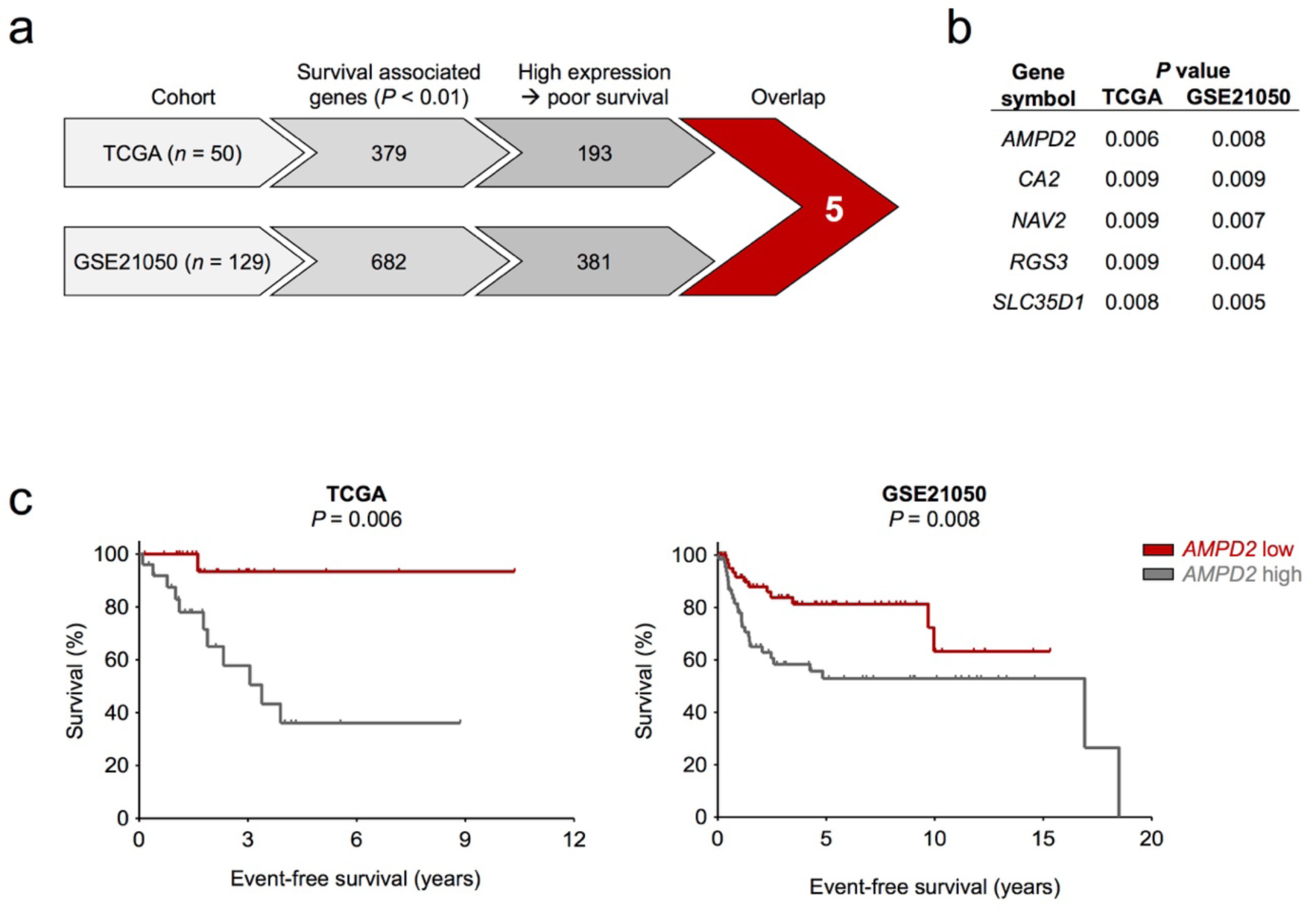
Subsequent filtering for survival associated genes in two independent RNA expression datasets for UPS reveals *AMPD2* as most promising candidate. a) Workflow of applied filtering steps in the survival analysis and number of genes meeting the criteria. b) List of the resulting candidates from Fig. 1a and their *P* values in both cohorts (Mantel-Haenszel test). c) Kaplan-Meier plots displaying the survival of patients with *AMPD2* low versus high expressing tumors in the TCGA and GSE21050 cohort (Mantel-Haenszel test).

### High AMPD2 protein levels correlate with worse outcome in UPS

As especially *AMPD2* appeared as an interesting candidate, we validated our findings based on the mRNA level by immunohistochemical staining for AMPD2 in a third UPS cohort comprising 83 samples (Table 1) on the protein level. To this end, we semi-quantitatively scored the intensity of AMPD2 immunoreactivity. In this cohort, 39 tumors exhibited no, 23 weak, 10 intermediate, and 11 strong AMPD2 immunoreactivity (Fig. 2a). For correlation with overall survival, samples were grouped in AMPD2-negative (corresponding to no immunoreactivity, *n* = 39), and AMPD2-positive (corresponding to weak to strong immunoreactivity, *n* = 44). Strikingly, detection of AMPD2 expression was associated with a significantly worse outcome, and early separation of the two groups in Kaplan-Meier analysis (Fig. 2b). Collectively, these findings made in three independent cohorts provide evidence that AMPD2 may constitute a general and robust prognostic marker in UPS.

**Figure 2.**
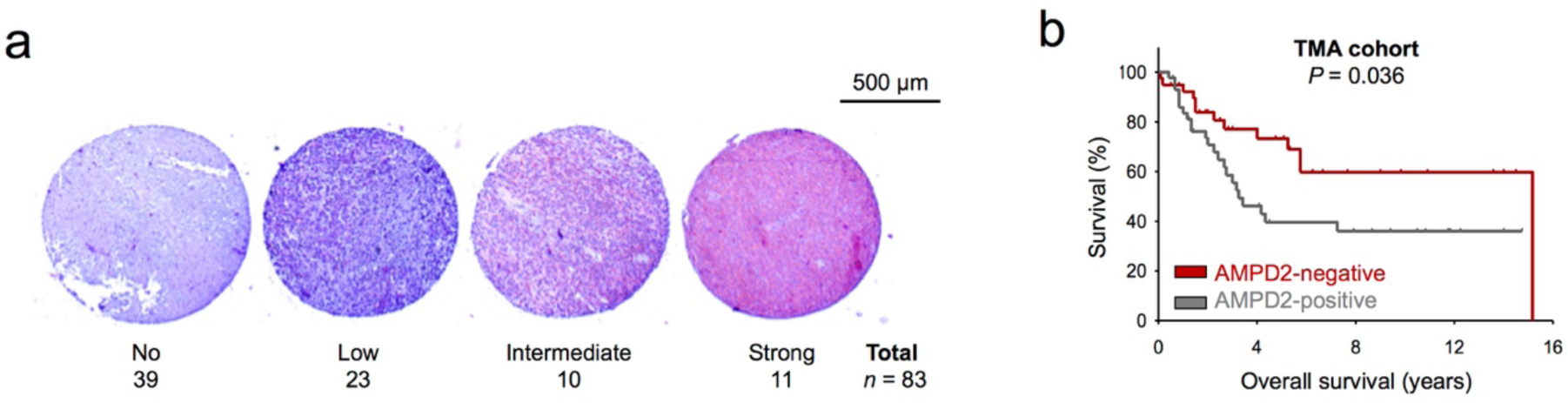
Staining of an UPS TMA for AMPD2 confirms its association with survival on protein level. a) Representative images of UPS tissue cores with no, low, intermediate, and strong cytosolic AMPD2 immunoreactivity. The number of individual UPS samples showing the corresponding staining intensities is reported. b) Kaplan-Meier plot displaying the overall survival of UPS patients carrying tumors without AMPD2 immunoreactivity (AMPD2-negative) and those with low to strong immunoreactivity (AMPD2-positive) (Mantel-Haenszel test).

### Copy number variations (CNVs) promote *AMPD2* overexpression in UPS

Since *AMPD2* appeared to be heterogeneously expressed in UPS, and since UPS typically feature high rates of genomic aberrations, we reasoned that CNVs at the *AMPD2* locus could cause the variable *AMPD2* expression. We therefore correlated CNV information and *AMPD2* expression data available for the TCGA cohort. In a linear regression analysis, we observed a highly significant correlation between the *AMPD2* expression levels and CNVs at the *AMPD2* locus in UPS (*r*_Pearson_ = 0.57; *P* < 0.001) (Fig. 3a).

**Figure 3.**
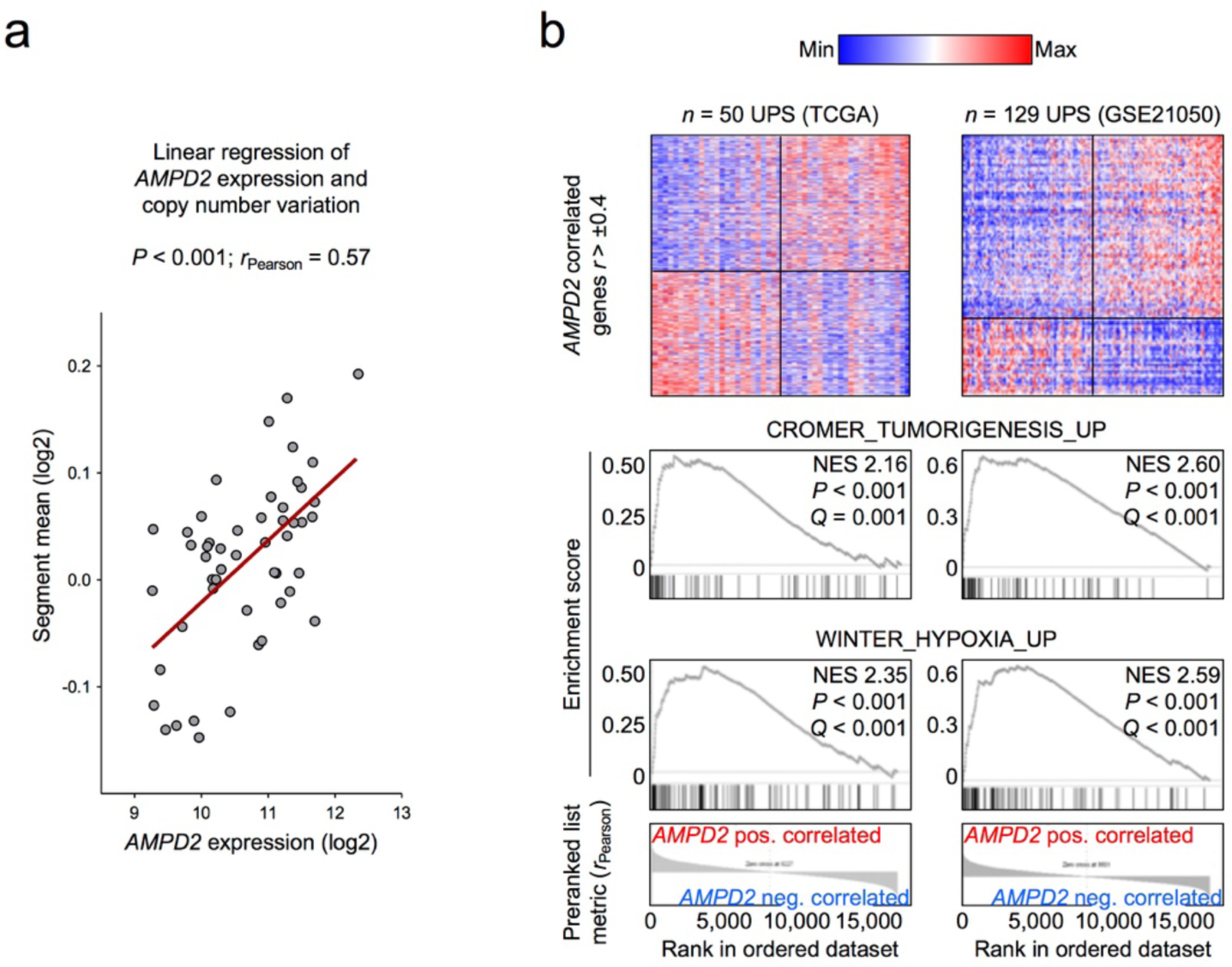
*AMPD2* expression correlates with copy number gains at the *AMPD2* locus and is associated with hypoxia and tumorigenesis. a) Dot-plot displaying the median *AMPD2* expression levels and copy number segment means at the *AMPD2* locus for each patient of the TCGA cohort. The red line represents the linear regression of the data, *r*_Pearson_ = 0.57, *P* < 0.001. b) Heat-maps for *AMPD2* co-and antiregulated genes in the TCGA and GSE21050 cohort with representative enriched gene sets within *AMPD2*-co-regulated genes. NES, normalized enrichment score.

**Figure 4.**
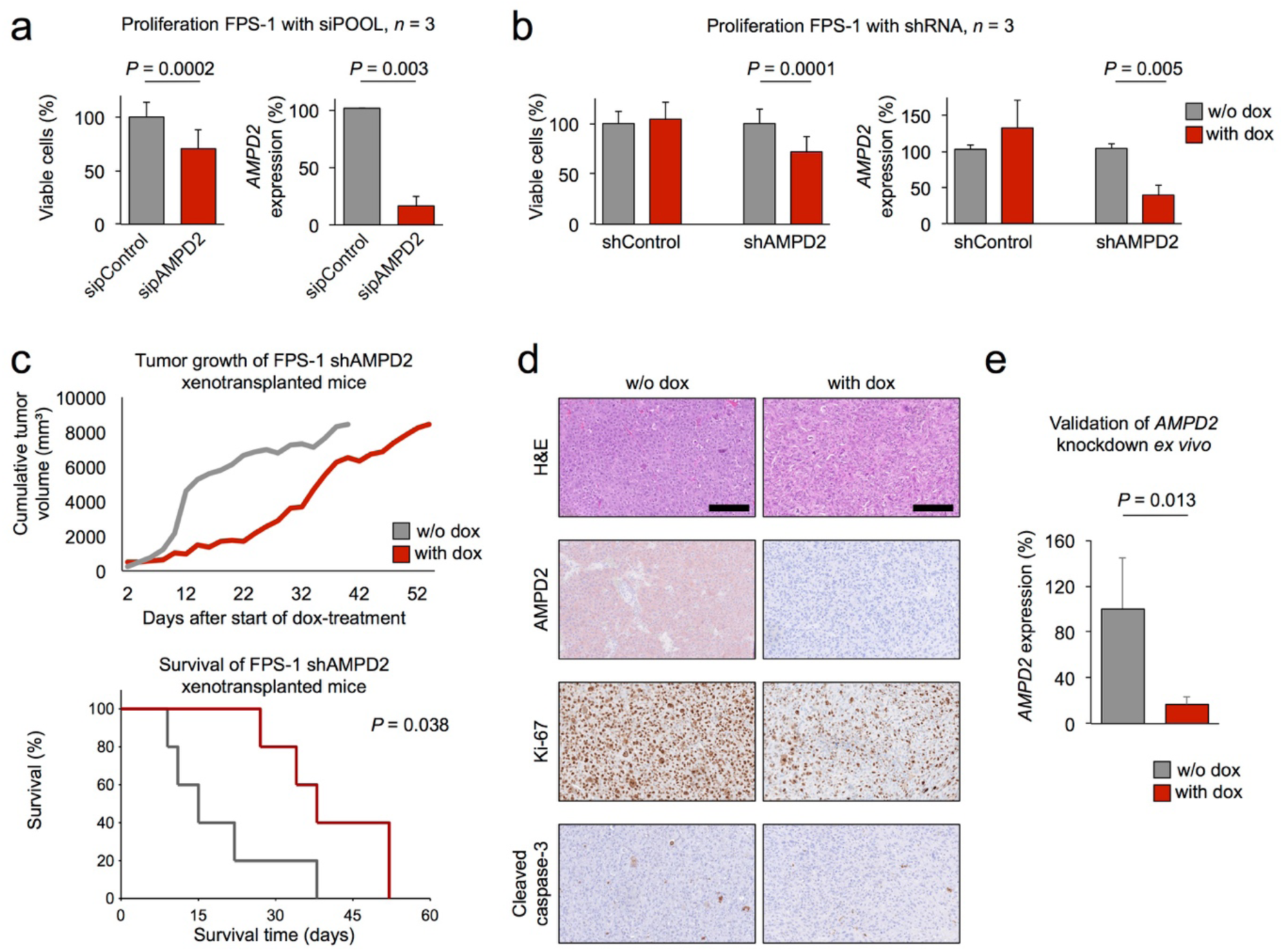
*AMPD2* silencing impairs UPS cell line proliferation *in vitro* and *in vivo*. a) Bar plots indicating viable cell count of sipControl (grey) and sipAMPD2 (red) transfected FPS-1 cells relative to control on the left, knockdown control on the right; whiskers indicate standard deviation. Statistical comparison by unpaired two-tailed student’s t-test. b) Left: bar plots indicating viable cell count of shControl and shAMPD2 transduced FPS-1 cells without (grey) and with (red) shRNA induction relative to control; right: knockdown control. Statistical comparison by unpaired two-tailed student’s t-test. c) Cumulative tumor volume over time for FPS-1 shAMPD2 xenotransplanted mice treated with doxycycline (dox; red) and controls (grey) and corresponding Kaplan-Meier plot below representing the survival time until average tumor diameter exceeded 15 mm (stop criterion) (Mantel-Haenszel test). d) Representative micrographs of the FPS-1 shAMPD2 tumors from mice without (left) and with (right) AMPD2 knockdown induction with dox (H&E, AMPD2, Ki-67, and cleaved caspase-3 staining; 20X magnification, scale bar = 200 µM). e) Bar plot for *AMPD2* knockdown validation for the FPS-1 shAMPD2 tumors from mice treated with dox in drinking water by qRT-PCR. Whiskers indicate standard deviation. Statistical comparison by unpaired two-tailed student’s t-test.

To get first clues on the potential biological relevance of CNV-mediated *AMPD2* overexpression in UPS, we performed gene-set enrichment analysis (GSEA) on *AMPD2* co-expressed genes in the TCGA and GSE21050 cohort. The gene-sets with highest normalized enrichment score (NES) were highly concordant in both cohorts (**Supplementary Tables 3 and 4**) and indicated that high *AMPD2* expression in UPS is associated with gene signatures involved in hypoxia and tumor growth (Fig. 3b).

### AMPD2 may promote proliferation and tumor growth of UPS

As our GSEA in primary tumors suggested that AMPD2 may have a functional role in growth of UPS, we aimed at validating this prediction in a cell line model. To this end, we took advantage of an established UPS cell line (FPS-1), which recapitulates key tumor features of UPS *in vivo*^15^ and performed knockdown experiments. In a first step, we serially transfected the FPS-1 cells with an siPOOL consisting of 30 specific siRNAs against *AMPD2* (sipAMPD2) and compared their proliferation with that of cells transfected with a non-targeting control siPOOL (sipControl). Transfection with the sipAMPD2 reduced *AMPD2* expression levels onto around 20% after 120 h, which was accompanied by a significant reduction of viable cells by around 30% (**Fig. 4a**). To validate this finding *in vivo*, we transduced FPS-1 cells with lentivirus containing a doxycycline (dox)-inducible short hairpin RNA (shRNA) against *AMPD2* (shAMPD2; pLKO-TET-ON all-in-one system^17^) or a non-targeting control shRNA (shControl). While dox-treatment had no effect on *AMPD2* expression and proliferation in FPS-1 shControl cells, treatment of FPS-1 shAMPD2 cells with dox significantly reduced *AMPD2* expression under 40% and efficiently impaired cell proliferation *in vitro* (Fig. 4b). We next injected FPS-1 shAMPD2 cells subcutaneously into the flanks of immunocompromised NSG mice and monitored tumor growth over time. Once most tumors reached an average diameter of 5 mm, we randomized the mice and henceforth treated half of them with dox in the drinking water (2 mg/dl). Strikingly, AMPD2 knockdown significantly delayed tumor growth (Fig. 4c), which was accompanied by a significant reduction in proliferating cells as indicated by IHC stain for Ki-67, while it did not induce apoptotic cell death as verified by staining for cleaved caspase-3 (Fig. 4d). The knockdown of AMPD2 was confirmed in each xenograft *ex vivo* by IHC and by qRT-PCR (Fig. 4d,e), which underscores the specificity of the used anti-AMPD2-antibody. Collectively, these results suggest that AMPD2 may promote proliferation and tumor growth of UPS.

## DISCUSSION

For many sarcoma subtypes including UPS, standard treatment regimens are often insufficient to achieve long-term disease control, resulting in local recurrence and/or metastasis, to which many patients will ultimately succumb^2,18–20^. To identify high-risk patients early on and to assign them upfront to adequate therapy regimens, novel prognostic and predictive biomarkers are urgently required. Although the diversity and rarity of sarcomas pose general obstacles for conducting statistically reliable biomarker studies in these cancers^21^, the integration of publicly available ‘omics’ data and functional assays might help overcoming this problem.

In the current study, we took a functional genomics approach by combining public gene expression datasets, immunohistochemistry, and cell-based *in vitro* and *in vivo* assays to identify *AMPD2* as a new prognostic candidate marker for UPS.

The relative small sample size of 50 in the TCGA cohort harbors the risk to miss genuine prognostic markers in survival analysis when correcting hundreds of potential candidates for multiple testing. We therefore consciously relinquished such statistical correction but chose instead, to only accept those genes as prognostic marker candidates with overlapping results in a second independent cohort. In these analyses, we focused on genes whose high expression correlated with worse outcome as they might also represent new therapeutic targets. While clinical TCGA data comprised the time of overall survival, the microarray dataset (GSE21050) contained information on metastasis-free survival. As metastasis is the predominant cause for disease specific death^22^, both datasets compared complimentary aspects of patient outcome. These analyses highlighted *AMPD2* as the most promising biomarker candidate, which was fully validated in a third independent cohort on the protein level.

AMPD2 is the liver isozyme of three known AMP deaminases, converting AMP to IMP, which is crucial for purine metabolism^10,23^. So far, the protein was reported to be linked with liver pathology^24,25^, but not with cancer^26^. In our FPS-1 UPS model, silencing of AMPD2 by RNA interference markedly reduced proliferation and tumor growth *in vitro* and *in vivo* without inducing apoptotic cell death. These findings suggest a functional relevance of AMPD2 in UPS, possibly through deregulation of the purine metabolism. Conversely, the frequently observed copy-number gains at the *AMPD2* locus and subsequent overexpression of AMPD2 in UPS primary tumors may facilitate tumor growth, and thus contribute to worse patient outcome. Future studies have to further characterize the functional role of AMPD2 in UPS, and to test whether it may additionally serve as a drug target for therapeutic blockage of increased purine metabolism. Also, additional work is necessary to validate the potential prognostic value of the other biomarker candidates that were identified in our transcriptome analyses (Fig. 1b).

Taken together, our data suggest that *AMPD2* is a promising novel prognostic biomarker for UPS. We therefore recommend its validation in prospective studies. Furthermore, our study highlights the importance of including even rare entities such as UPS in ongoing cancer genomics projects such as TCGA, and our functional genomics approach may serve as a blueprint for identification and validation of additional biomarkers for other rare cancers.

## Abbreviations

CDF: chip description file
dox: doxycycline
FTPE: formalin fixed paraffin embedded
GEO: gene expression omnibus
GSEA: gene set enrichment analysis
H&E: hematoxylin and eosin
NES: normalized enrichment score
NSG: NOD/Scid/gamma
qRT-PCR: quantitative real time polymerase chain reaction
TCGA: The Cancer Genome Atlas
TMA: tissue microarray
UPS: undifferentiated pleomorphic sarcoma

## Author contributions

M.F.O. and T.P.G.P. conceived the study, wrote the paper and designed the figures. M.F.O. performed bioinformatic analyses and experiments. J.S.G. programmed the in-house survival analysis tool. T.Kn. scored IHC staining of TMAs and A.A.H. correlated the results with clinical data. J.M., R.A.R., S.S., G.S., S.O., J.L, M.D., M.C.B., and A.M. supported the experimental procedures. M.H. provided the FPS-1 cell line. T.Ki. provided lab infrastructure and histological guidance. E.B., T.D., and T.P.G.P. supervised the project and provided scientific advice. All authors read and approved the final version of this manuscript.

## ACKNOWLEDGEMENTS

We thank Mrs. Andrea Sendelhofert and Mrs. Anja Heier for excellent technical assistance.

## REFERENCES

1 Fletcher DM, Bridge JA, Hogendoorn PCW, et al. WHO Classification of Tumours of Soft Tissue and Bone, 4th Edition. Lyon, France: International Agency of Research on Cancer, 2013.

2 Fairweather M, Keung E, Raut CP. Neoadjuvant Therapy for Soft-Tissue Sarcomas. Oncol Williston Park N 2016;30:99–106.

3 Matushansky I, Taub RN. Adjuvant chemotherapy in 2011 for patients with soft-tissue sarcoma. Nat Rev Clin Oncol 2011;8:434–438.

4 Mariño-Enríquez A, Bovée JVMG. Molecular Pathogenesis, Diagnostic, Prognostic and Predictive Molecular Markers in Sarcoma. Surg Pathol Clin 2016;9:457–473.

5 Cancer Genome Atlas Research Network. Electronic address, Cancer Genome Atlas Research Network. Comprehensive and Integrated Genomic Characterization of Adult Soft Tissue Sarcomas. Cell 2017;171:950–965.e28.

6 Savina M, Le Cesne A, Blay J-Y, et al. Patterns of care and outcomes of patients with METAstatic soft tissue SARComa in a real-life setting: the METASARC observational study. BMC Med 2017;15:78.

7 Nascimento AF, Raut CP. Diagnosis and management of pleomorphic sarcomas (so-called “MFH”) in adults. J Surg Oncol 2008;97:330–339.

8 Delisca GO, Mesko NW, Alamanda VK, et al. MFH and high-grade undifferentiated pleomorphic sarcoma-what’s in a name? J Surg Oncol 2015;111:173–177.

9 Widemann BC, Italiano A. Biology and Management of Undifferentiated Pleomorphic Sarcoma, Myxofibrosarcoma, and Malignant Peripheral Nerve Sheath Tumors: State of the Art and Perspectives. J Clin Oncol Off J Am Soc Clin Oncol 2018;36:160–167.

10 Van den Bergh F, Sabina RL. Characterization of human AMP deaminase 2 (AMPD2) gene expression reveals alternative transcripts encoding variable N-terminal extensions of isoform L. Biochem J 1995;312 (Pt 2):401–410.

11 Chibon F, Lagarde P, Salas S, et al. Validated prediction of clinical outcome in sarcomas and multiple types of cancer on the basis of a gene expression signature related to genome complexity. Nat Med 2010;16:781–787.

12 Irizarry RA, Hobbs B, Collin F, et al. Exploration, normalization, and summaries of high density oligonucleotide array probe level data. Biostat Oxf Engl 2003;4:249–264.

13 Dai M, Wang P, Boyd AD, et al. Evolving gene/transcript definitions significantly alter the interpretation of GeneChip data. Nucleic Acids Res 2005;33:e175.

14 Subramanian A, Tamayo P, Mootha VK, et al. Gene set enrichment analysis: a knowledge-based approach for interpreting genome-wide expression profiles. Proc Natl Acad Sci U S A 2005;102:15545–15550.

15 Hakozaki M, Hojo H, Sato M, et al. Establishment and characterization of a new cell line, FPS-1, derived from human undifferentiated pleomorphic sarcoma, overexpressing epidermal growth factor receptor and cyclooxygenase-2. Anticancer Res 2006;26:3393–3401.

16 Hannus M, Beitzinger M, Engelmann JC, et al. siPools: highly complex but accurately defined siRNA pools eliminate off-target effects. Nucleic Acids Res 2014;42:8049–8061.

17 Wiederschain D, Wee S, Chen L, et al. Single-vector inducible lentiviral RNAi system for oncology target validation. Cell Cycle Georget Tex 2009;8:498–504.

18 Rothermundt C, Whelan JS, Dileo P, et al. What is the role of routine follow-up for localised limb soft tissue sarcomas? A retrospective analysis of 174 patients. Br J Cancer 2014;110:2420–2426.

19 Kang S, Kim H-S, Kim S, et al. Post-metastasis survival in extremity soft tissue sarcoma: A recursive partitioning analysis of prognostic factors. Eur J Cancer 2014;50:1649–1656.

20 Stiller CA, Trama A, Serraino D, et al. Descriptive epidemiology of sarcomas in Europe: Report from the RARECARE project. Eur J Cancer 2013;49:684–695.

21 Brennan MF, Antonescu CR, Moraco N, et al. Lessons learned from the study of 10,000 patients with soft tissue sarcoma. Ann Surg 2014;260:416–422.

22 Mehlen P, Puisieux A. Metastasis: a question of life or death. Nat Rev Cancer 2006;6:449–458.

23 Mahnke-Zizelman DK, van den Bergh F, Bausch-Jurken MT, et al. Cloning, sequence and characterization of the human AMPD2 gene: evidence for transcriptional regulation by two closely spaced promoters. Biochim Biophys Acta 1996;1308:122–132.

24 Lanaspa MA, Cicerchi C, Garcia G, et al. Counteracting roles of AMP deaminase and AMP kinase in the development of fatty liver. PloS One 2012;7:e48801.

25 Cicerchi C, Li N, Kratzer J, et al. Uric acid-dependent inhibition of AMP kinase induces hepatic glucose production in diabetes and starvation: evolutionary implications of the uricase loss in hominids. FASEB J Off Publ Fed Am Soc Exp Biol 2014;28:3339–3350.

26 Szydłowska M, Roszkowska A. Expression patterns of AMP-deaminase isozymes in human hepatocellular carcinoma (HCC). Mol Cell Biochem 2008;318:1–5.

